# Exploration of Natural Products for Targeting IDH1/2 Mutations in Acute Myeloid Leukemia through Ligand-Based Pharmacophore Screening, Docking, ADME-T and Molecular Dynamic Simulation Approaches

**DOI:** 10.1101/2024.08.27.609840

**Authors:** Uddalak Das, Dheemanth Reddy Regati, Jitendra Kumar, R. Sowdhamini

## Abstract

**Background:** Mutations in isocitrate dehydrogenase 1 (IDH1) and 2 (IDH2) are prevalent drivers of acute myeloid leukemia (AML). While targeted therapies exist, resistance can emerge. This study explored the potential of natural products to identify novel dual IDH inhibitors.

**Methods:** *In-silico* screening of the COCONUT database was performed using Lipinski’s Rule of Five. Pharmacophore modelling identified crucial features for IDH binding. Docking simulations with Glide (Schrödinger) assessed binding affinity, followed by MM-GBSA calculations for free energy estimation. The most promising candidate underwent ADME/T and toxicity analysis. Finally, molecular dynamics (MD) simulations evaluated the stability of protein-ligand complexes and binding interactions, followed by trajectory analysis using DCCM and PCA.

**Results:** The study identified Ternstroside D (CNP0166496) as a potential dual inhibitor for IDH1 and IDH2 mutations. Docking simulations and MM-GBSA calculations predicted favourable binding affinities. MD simulations revealed stable protein-ligand complexes, and in-silico ADME/T analysis suggested good drug-like properties and a favorable safety profile for CNP0166496.

**Conclusion:** This *in-silico* study provides compelling evidence for Ternstroside D (CNP0166496) as a promising dual inhibitor for IDH1 and IDH2 mutations in AML. Further *in-vitro* and *in-vivo* studies are warranted to validate these findings.

## 1. INTRODUCTION

Acute myeloid leukemia (AML) is a malignant disorder that targets hematopoietic stem cells or progenitor cells, characterized by an excessive proliferation of myeloblasts. This results in a reduction of terminally differentiated cells such as white blood cells, platelets, and red blood cells, leading to severe health implications (El-Jawahri *et al*., 2019). The American Cancer Society projected approximately 60,650 new cases of leukemia in the US in 2022, with AML being the most lethal subtype, accounting for 48% of leukemia-related deaths (Miller *et al*., 2022). Despite chemotherapy being the first-line treatment, the five-year survival rate for AML patients remains low at about 28%, with an overall median survival of only 5 to 10 months in older patients or those with comorbidities (Miyamoto *et al*., 2022).

AML is often associated with the dysregulated expression of certain genes, including the isocitrate dehydrogenase (IDH) genes, which play a crucial role in the disease’s pathogenesis. Mutations in IDH1 and IDH2 were first identified in AML in 2008 and are present in about 20% of cases (Issa & DiNardo, 2021). These mutations occur at conserved arginine residues within the enzyme’s active site, specifically at R132 for IDH1 and at R140 or R172 for IDH2 (Zarnegar-Lumley *et al*., 2023). Mutant IDH1 and IDH2 enzymes alter the Krebs cycle and convert isocitrate to 2-hydroxyglutarate (2-HG) instead of alpha-ketoglutarate, leading to the accumulation of 2-HG, which acts as an oncometabolite and disrupts cellular differentiation and proliferation (Han *et al*., 2020). However, there are no reported cancer associated mutations in IDH3(Liu *et al*., 2023). 2-HG inhibits α-KG-dependent enzymes like TET and lysine demethylases, acting as an oncogenic metabolite and causing a hyper-methylated phenotype. Its accumulation leads to abnormal histone and DNA methylation, blocks cell differentiation, and promotes leukemogenesis (Du & Hu, 2021). Thus, targeting mutant IDH enzymes is crucial for AML therapy. The development of IDH inhibitors has been a significant advancement in AML therapy. The FDA has approved several IDH inhibitors based on clinical trials demonstrating promising responses. However, the efficacy of these inhibitors compared to standard treatments remains under debate (Fathi *et al*., 2020). Consequently, there is an urgent need for individualized treatments with higher efficacy and better safety profiles.

Natural products have been a valuable source of therapeutic agents, with many anticancer drugs being derived from natural sources (Cragg & Pezzuto, 2016). This study screens and investigates the potential of 4,07,270 bioactive compounds from COCONUT database as dual inhibitors of IDH1_R132H and IDH2_R140Q for treating AML using advanced *in silico* methods such as structure-based virtual screening, molecular docking, and molecular dynamics (MD) simulations have become invaluable tools in drug discovery. These computational approaches are cost-effective and time-efficient compared to traditional experimental methods (Sadybekov & Katritch, 2023). In this study, we employed a combination of diverse ligand-based pharmacophore modelling, molecular docking, and MMGBSA to screen and validate potential IDH dual inhibitors from the CONONUT database. Additionally, we performed ADMET profiling and molecular dynamics simulations to assess the stability and drug-likeness of the identified compounds.

Our integrated approach aims to identify potent dual inhibitors of IDH1 and IDH2 mutations, contributing to the development of more effective treatments for AML. By leveraging natural products and computational techniques, this research provides valuable insights into the discovery of novel therapeutic agents for leukemia and other IDH-related malignancies.

## 2. MATERIALS AND METHODS

### 2.1. Dataset generation and screening

The COCONUT (version 2022, Collection of Open Natural ProdUcTs Online) database was utilized as the screening source, containing approximately 407,270 molecules (Sorokina *et al*., 2021). The molecules were then filtered using QikProp module in “Structure Filtering” option Schrödinger 2023-1, with zero violation of Lipinski’s Rule of 5 generating an output of 276,409 molecules. The filtered structures were then prepared using the LigPrep Module with the OPLS4 force field (Lu *et al*., 2021). The possible states of molecules were generated using Epik [49] at pH 7.0 ± 2.0, retained specified chirality, followed by tautomer generation and the generation of up to 16 low-energy conformations per ligand. The total number of generated structures were around 3.57 million.

### 2.2. Pharmacophore modelling/screening

#### 2.2.1. Pharmacophore hypothesis generation

A pharmacophore comprises distinct characteristics like steric properties, hydrogen bond donor (HBD), hydrogen bond acceptor (HBA), and electronic chemical attributes. These features indicate how a compound functions uniquely within the active biological site (S.-Y. Yang, 2010). The development of ligand-based pharmacophore design relied on identified actives with confirmed pharmacological effects targeting the chosen receptor.

To generate the pharmacophore model, an initial subset comprising the first 100 compounds linked to the IDH1 and IDH2 was extracted from the binding database, and sorted based on their respective IC_50_ values. Three approved drugs that are effective in IDH1 inhibition, namely *Enasidenib* (89683805), *Olutasidenib* (118955396), and *Ivosidenib* (71657455) was used along with the relevant conformers for benchmark datasets for IDH1 and *Enasidenib* (89683805) for IDH2 (Ganji *et al*., 2023). This curated dataset was the foundation for subsequent pharmacophore model construction and screening. Within the LigPrep module, the OPLS4 force field was engaged (Lu *et al*., 2021), concomitant with default parameters for ionization. Furthermore, routine procedures encompassing desalination, tautomerization, and computational adjustments were implemented per software defaults. High-quality 3D structures for the drug-like molecules were prepared.

In the Develop Pharmacophore Model module, the hypothesis match was set to 75% for both and the number of features in the hypothesis was kept from 4 to 7 with the preferred number to 5. The ranking and scoring of the hypothesis were set to the default “Phase Hypo Score” (Yu *et al*., 2021).The Generate Conformer and Minimize Output conformer options were activated, with the target number of conformers set to 50 (Cole *et al*., 2018).

#### 2.2.2. Validation of the pharmacophore model

The developed pharmacophore model was verified to check its ability to predict the activity of new compounds effectively. This procedure is a prerequisite before the pharmacophore model can be employed for virtual screening (Kaserer *et al*., 2015). The parameters used for evaluating the efficiency of the developed pharmacophore model are enrichment factor (EF), receiver operating characteristic (ROC) curves, Boltzmann-enhanced discrimination of ROC (BEDROC), and robust initial enhancement (RIE) (Truchon & Bayly, 2007).

A decoy set was created using the Generate DUDE Decoys program, which is found at http://dude.docking.org/generate (Mysinger *et al*., 2012). For converting the output into 3D structures, Open Babel v2.4.1 was used (O’Boyle *et al*., 2011). Ligand preparation was done following the default settings and the protocols of LigPrep. The OPLS4 force field (Lu *et al*., 2021) has been employed in the minimization procedure.

#### 2.2.3. Screening using Pharmacophore Model

The prepared ligands were screened using the best generated pharmacophore model, DDHR_1 for IDH1 and ADDHR_1 for IDH2 using the “Phase Module”. Prefer partial matches of more features were activated. All the pharmacophore properties (4 of 4 for IDH1 and 5 of 5 for IDH2) were selected for screening. The pharmacophore model screened ∼3.57 million structures to ∼181K structures and ∼67K for IDH1 and IDH2 respectively.

### 2.3. Molecular Interactions

#### 2.3.1. Protein preparation

The X-ray crystallographic structure of the IDH1 and IDH2 target protein with PDB ID: 5DE1 and 5I96, found at https://www.rcsb.org, with a resolution of 2.25Å and 1.55Å respectively, by Yen *et. al.* (2017) (Yen *et al*., 2017), was used in the study. The structure analysis was carried out using the PDBsum web server (https://www.ebi.ac.uk/thornton-srv/databases/pdbsum) (Laskowski *et al*., 2018). PDBsum online server was also used to check the validation of the IDH1/2 with the Ramachandran plot.

In the Schrödinger Maestro protein preparation wizard, the proteins was pre-processed with the PROPKA module for an optimization of H-bonds (Gokcan & Isayev, 2022), followed by minimization of structures towards convergence of heavy atoms at RMSD 0.3Å using OPLS4 force field and removal of water molecules more than 5Å away from ligands afterward (Lu *et al*., 2021).

#### 2.3.2. Receptor grid generation

The receptor grid was generated keeping the hydrophobic region and the region where co-crystallized inhibitor GSK321A and AG-221 (Enasidenib) for IDH1 and IDH2 respectively was attached to the complex, at the centroid of the grid. The coordinates of the receptor grid for IDH1 were X= −21.12, Y = 11.78, Z= 6.25 and for IDH2 were X=-1.53, Y=15.61, Z= - 24.69.

#### 2.3.3. Molecular docking

Docking was limited to ligands with 100 rotatable bonds and fewer than 500 atoms. Van der Waals radii scaling factor was set to 0.80, with a partial charge cutoff of 0.15 (Pavlin *et al*., 2019). Sample nitrogen inversions and sample ring conformations were activated, and the ligand sampling was set to flexible. All predicted functional groups had bias sampling of torsions enabled. The module was configured to promote intramolecular hydrogen bonds and improve conjugated pi groups’ planarity.

Docking was done through a series of hierarchical filters i.e. HTVS mode (high-throughput virtual screening) for efficiently screening million compound libraries, to the SP mode (standard precision) for reliably docking tens to hundreds of thousands of ligands with high accuracy, to the XP mode (extra precision) where further elimination of false positives is accomplished by more extensive sampling and advanced scoring, resulting in even higher enrichment. Each step proceeded with the top 10% from the previous one (Bagchi *et al*., 2017). The HTVS filtered the library to 1,40,350, SP filtered to 25,872 and, XP filtered to around 2,589 structures for IDH1 and 4,865, 475, and 49 respectively for IDH2 (Friesner *et al*., 2004, 2006; Halgren *et al*., 2004; Y. Yang *et al*., 2021).

#### 2.3.4. MM-GBSA Screening

The prime module was utilized to predict the energy parameters obtained from the MM-GBSA (Molecular Mechanics-Generalized Born Surface Area) simulation. It aimed to predict the amounts of the stabilization energy coming from the potential interaction between the selected ligands and the target receptors 5DE1 and 5I96 (Genheden & Ryde, 2015). The VSGB solvation model was used and the force field was set as OPLS4 (Li *et al*., 2011).

The top 10% of the ligands generated through XP docking were analysed using MM-GBSA for IDH1 and all the compounds generated through XP for IDH2. And again, the compound having binding affinity with both the proteins was selected from the top 10% of MMGBSA results. The only compound that matched the criteria was CNP0166496 (Ternstroside D) which was used for further ADME-Tox and MD Simulation studies.

### 2.4. *In silico* ADME/T and toxicity analysis

QikProp Module of Schrödinger, SwissADME (Daina *et al*., 2017), ProTox-3.0 (Banerjee *et al*., 2024), ADMETlab 2.0 (Xiong *et al*., 2021), and Deep-PK (Pires *et al*., 2015) server were used in the analysis of pharmacokinetic properties to assess the detailed ADMET properties of CNP0166496 (Ternstroside D).

### 2.5. Molecular Dynamics (MD) simulation studies

Desmond package was used to carry out the molecular dynamics simulations for the 5DE1-CNP0166496 and 5I96-CNP0166496 complexes. Each system was placed individually in an orthorhombic water box of 10 Å using the TIP3P water model (Mark & Nilsson, 2001). The ligand-protein complexes were modelled by the OPLS4 force field (Lu *et al*., 2021). Counter ions (Na+) were introduced in the ligand-protein complex structures to neutralize the total charge of the systems undergoing MD simulation. Furthermore, the energies of the systems were minimized to a minimum level using 2000 steps before initiating the MD simulation along an NPT lattice trajectory (Al-Jumaili *et al*., 2023).

The RMSD primarily suggests the stability of the ligand interaction, while RMSF describes the fluctuation and flexibility of the residues within the protein, particularly within the active site that is crucial for drug discovery. RMSD is calculated by the discovery 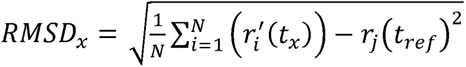. RMSF values also help in defining protein structures as they provide information about local conformational changes within the protein chains. The RMSF values are in units of Å and is calculated by the following equation 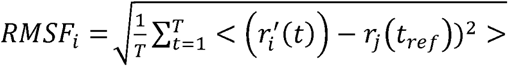 (Ghahremanian *et al*., 2022).

### 2.6. Principal Component Analysis (PCA) and Dynamical Cross-Correlation Matrix (DCCM) Analysis

The trajectory file generated after the MD simulation was converted to .dcd format with the help of VMD 1.9.3 software (Humphrey *et al*., 1996), which was subsequently uploaded to the MDM-TASK web server to perform PCA and DCCM analysis (Amamuddy *et al*., 2021). PCA was conducted to identify the major conformational changes and essential dynamics of the protein-ligand complex, while DCCM analysis was used to examine the correlated motions of residues over the simulation period.

### 2.7. MM-GBSA Analysis

The MM-GBSA-based binding free energy calculations were done on the 100ns long MD trajectories. For the selection of protein-ligand complexes, the binding energies calculated by this approach are more efficient than the glide score values. The main energy elements like H-bond interaction energy (ΔG_Bind_Hbond_), electrostatic solvation free energy (ΔG_Bind_Solv_), Coulomb or electrostatics interaction energy (ΔG_Bind_Coul_), lipophilic interaction energy (ΔG_Bind_Lipo_), and van der Waals interaction energy (ΔG_Bind_vdW_) altogether were considered to the calculation of MM-GBSA based relative binding affinity.

## 3. RESULTS

### 3.1. The IDH Proteins

Analysis through PDBsum web server identified 18 α-helices, 3 β-hairpins, 2 β-bulges, 17 β-turns as well as 20 helix-helix interactions for 5DE1 (IDH1) and 18, 3, 1, 18 and 25 respectively for 5I96 (IDH2).

Furthermore, the Ramachandran plot was also used to validate proteins, and it indicated that 92% of the residues were in preferred regions, 7.4% were in additional residue regions, 0.6% in generous regions, and 0.0% in disallowed regions. and 91.7%, 7.3%, 1.0% and 0.0% for IDH2. The overall G-Factor was 0.26 for IDH1 and 0.15 for IDH2. (Ramachandran *et. al.,* 1963) (see Figure 2)

**Figure 1.**
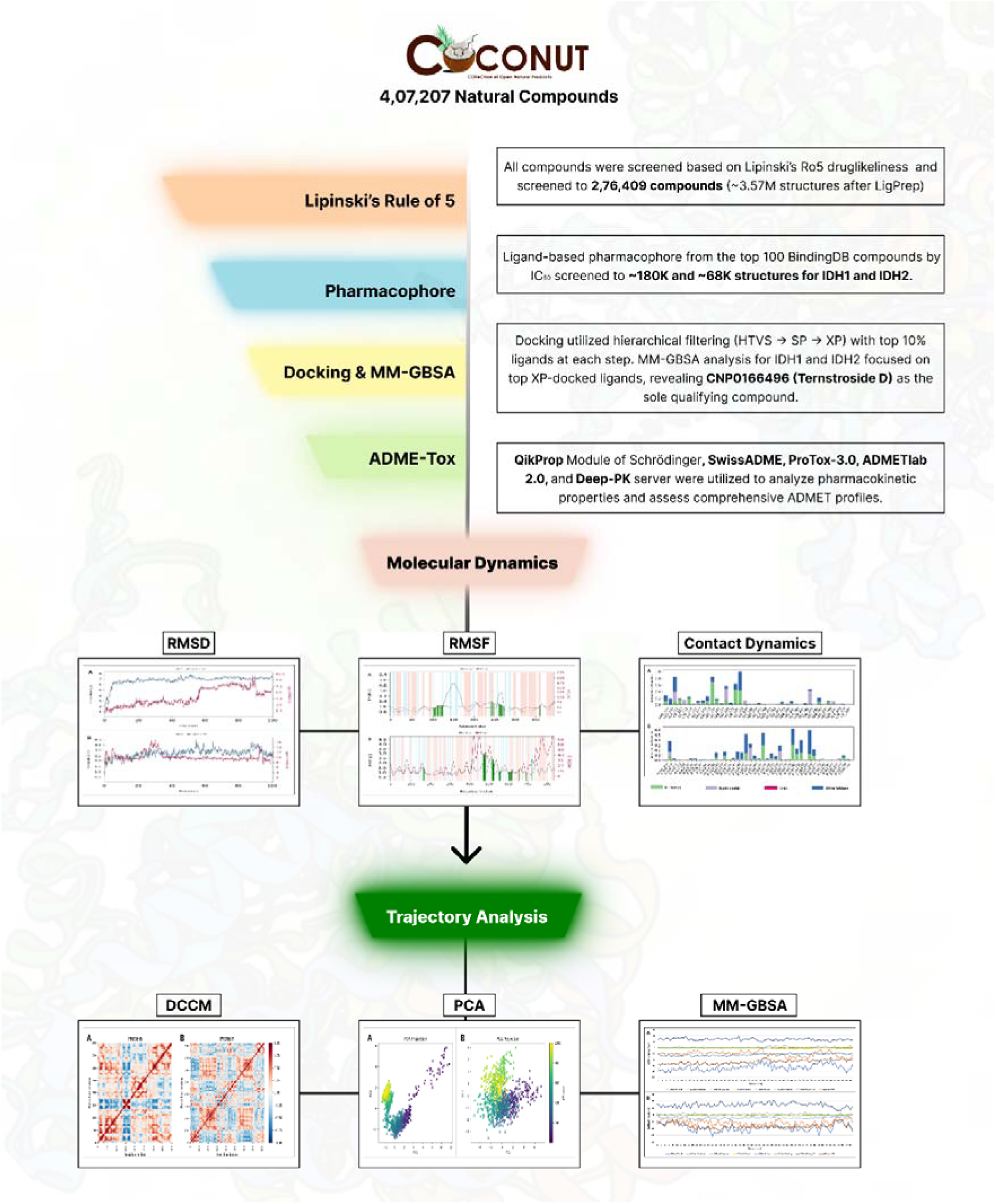
(Graphical Abstract) The schematic diagram of the complete dataset screening steps used in this study

**Figure 2.**
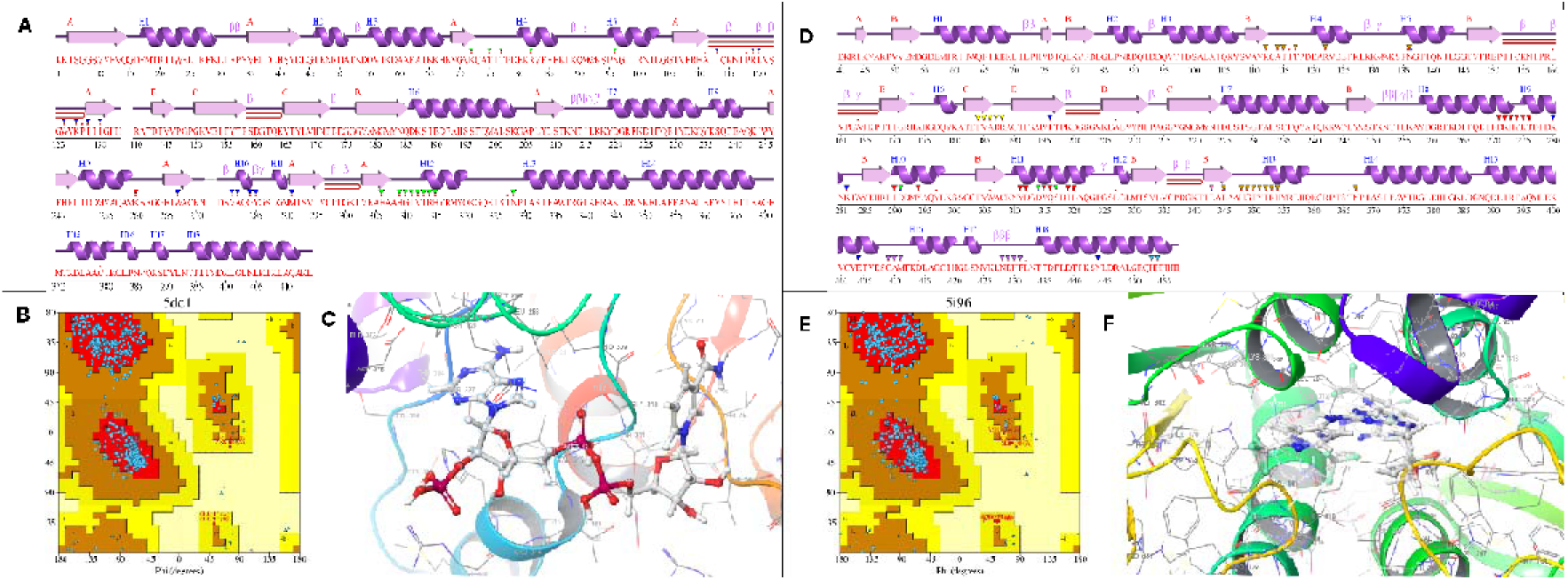
A: The secondary structure of the amino acid sequence of protein 5DE (generated by PDBsum); B: Ramachandran plot of the protein. The colour red indicates Iow-energy regions, yellow - allowed regions, pale yellow - generously allowed regions, and white - disallowed regions. C: The 3D structure of the IDH1 (Chain B) protein where the co-crystallized inhibitor GSK321A interacts; D: The secondary structure of the amino acid sequence of protein 5I96 (generated by PDBsum); B: Ramachandran plot of the protein. C: The 3D structure of the IDH2 protein where the co-crystallized inhibitor AG-221 (Enasidenib) interacts

**Figure 3.**
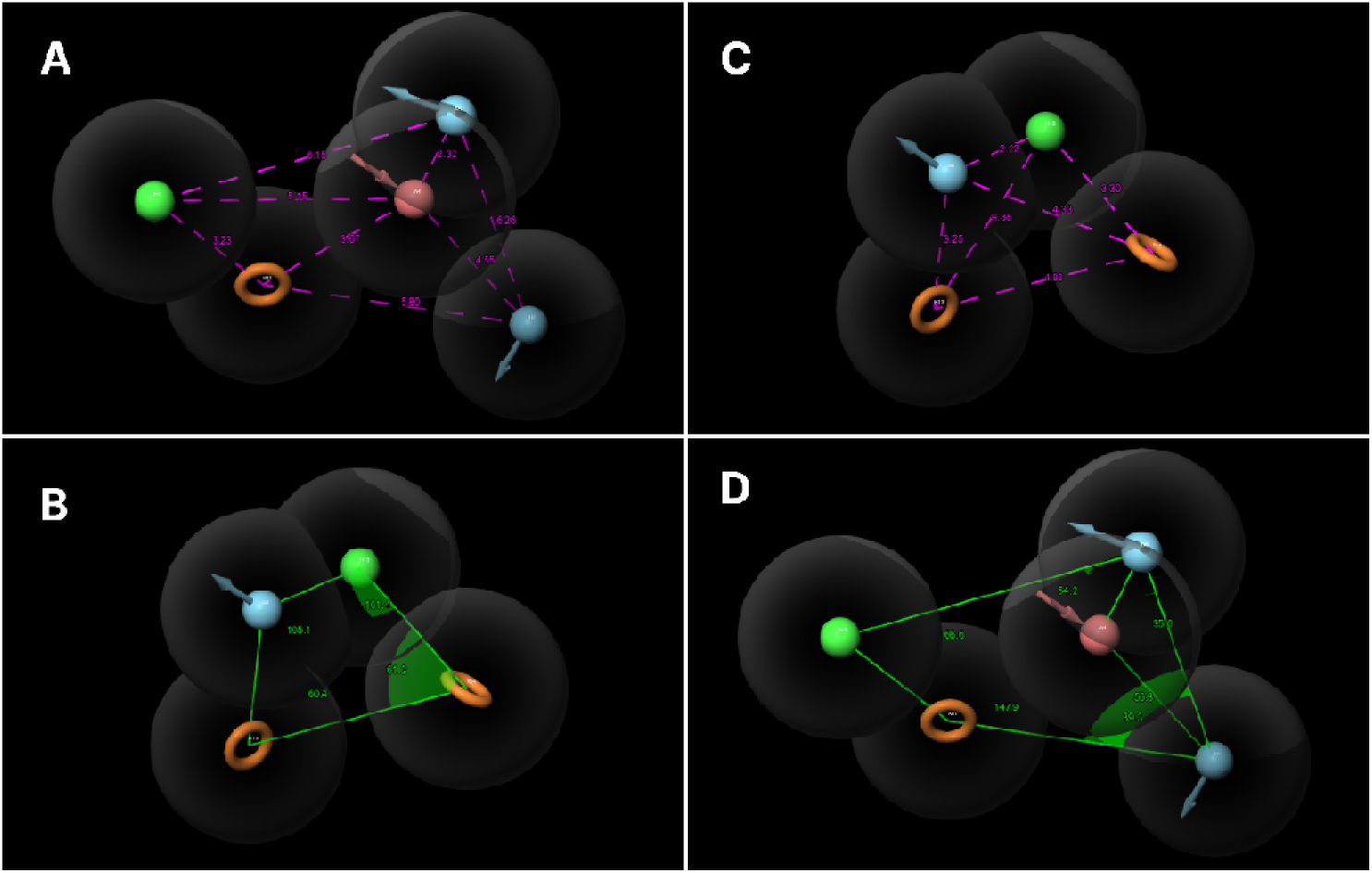
The distance (A) and angles (B) between pharmacophore features of the best pharmacophore model, DHRR_1 for IDH1 and ADDHR_1 (C and D) for IDH2

**Figure 4.**
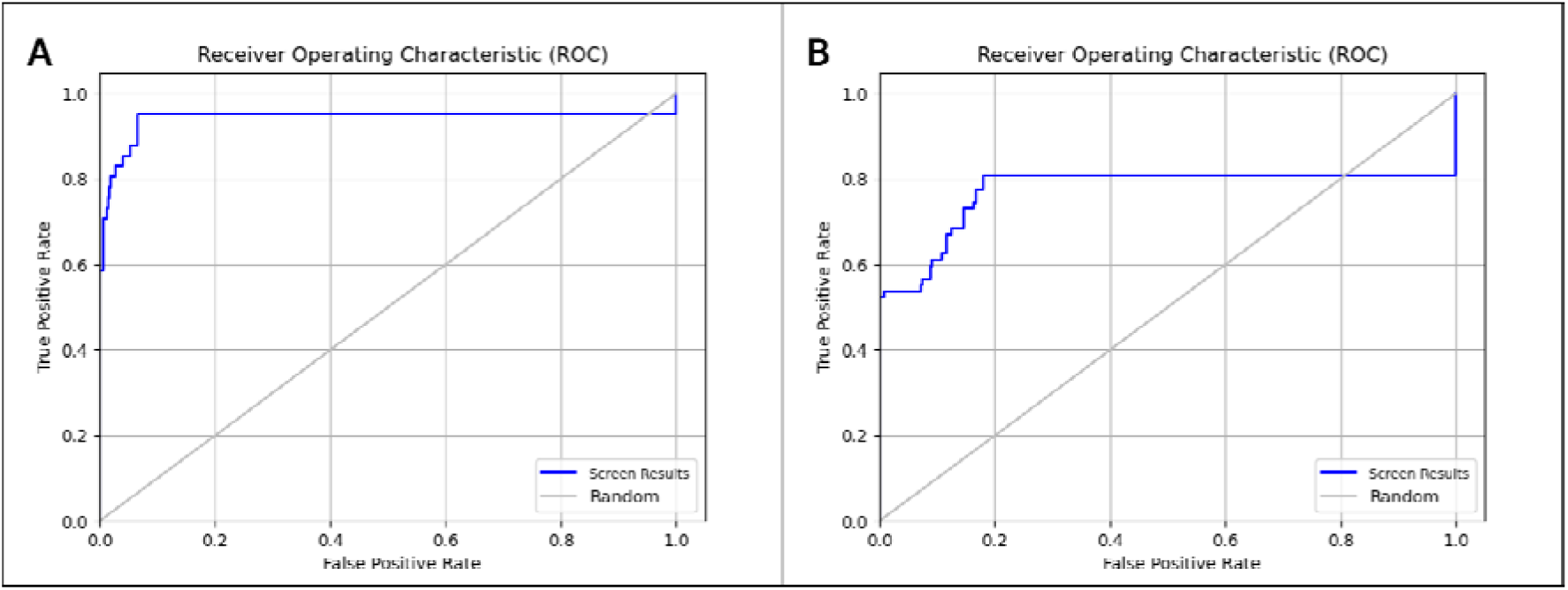
ROC curve represents the pharmacophore validation of the model A: DHRR_1 and B: ADDHR_1

### 3.2. Pharmacophore modelling/screening

The generated pharmacophore models were ranked automatically based on the survivability and BEDROC Score. For, IDH1, the hypothesis DHRR_1 revealed the highest Phase Hypo Score of 1.328 comprising one hydrogen bond donor (D), one hydrophobic (H) and two aromatic ring (R) features. The survival score of the hypothesis is 6.078, site score is 0.779, vector score is 0.933, volume score is 0.634, selectivity score is 1.449, and BEDROC score is 0.963. For IDH2, the hypothesis ADDHR_1 revealed the highest PhaseHypo Score of 1.330 comprising one hydrogen bond acceptors (A), two hydrogen bond doner (D), one hydrophobic (H), and one aromatic ring (R) features. The survival score of the hypothesis is 6.248, site score is 0.837, vector score is 0.943, volume score is 0.723, selectivity score is 1.703, and BEDROC score is 0.956.

The ∼3.5 million prepared ligand structures were screened using the DHRR_1 pharmacophore model and screened to 1,81,118 structures for IDH1 and to 67,859 structures using ADDHR_1 for IDH2.

### 3.3. Pharmacophore Models Validation

The pharmacophore model developed was rigorously validated to assess its ability to accurately predict the activity of novel compounds identified through database screening or synthesized de novo. Validation is an essential step before utilizing a pharmacophore model for virtual screening (Kaserer *et al*., 2015).

The DHRR_1 model was applied to a test set database comprising 200 inactive molecules (551 structures after preparation) generated via the DUDE Decoys tool. It screened the decoy set to only 35 structures indicating an efficiency of 93.648%. Further, validation of the DHRR_1 hypothesis revealed that EF in the top 1% of the decoy dataset is 15.93%, demonstrating that pharmacophore model is 15.93-fold efficient in detecting true positives/actives from the entire dataset. ROC score, RIE, and AUAC values were calculated as 0.77, 8, and 0.82, respectively. Thus, DHRR_1 was statistically significant in picking the actives from the decoy dataset. Statistical significance of model was also validated by calculating BEDROC. Contrary to EF, BEDROC seeks to measure the early enrichment of the actives. BEDROC values were calculated at different tuning parameter values (α□=□8.0, α□=□20.0, and α□=□160.9) and found to be 0.686, 0.702, and 0.995, respectively.

Similarly, the ADDHR_1 model was used to screen the same dataset, screening to 37 structures indicating 93.285% efficiency. The validation of the ADDHR_1 hypothesis showed that the enrichment factor (EF) in the top 1% of the decoy dataset is 25.39%, indicating that the model is 25.39 times more effective in identifying true positives/actives from the entire dataset. The ROC score, RIE, and AUAC values were found to be 0.94, 12.04, and 0.94, respectively, confirming that ADDHR_1 is statistically significant in selecting actives from the decoy dataset. The model’s statistical significance was further confirmed by calculating BEDROC, which, unlike EF, focuses on the early enrichment of actives. BEDROC values were computed at different tuning parameters (α□=□8.0, α□=□20.0, and α□=□160.9) and were determined to be 0.900, 0.870, and 0.983, respectively.

### 3.4. Binding Affinity Predictions

The docking score range for the top 500 hits compounds was found between −16.131 to - 11.628 kcal/mol for IDH1, after XP docking, and-16.822 to −7.017 kcal/mol for the 49 compounds for IDH2. The compound having binding affinity with both the proteins was in the top MMGBSA results was CNP0166496 (Ternstroside D) with ΔG_bind_ energy of 84.45 kcal/mol and −60.73 kcal/mol for IDH1 and IDH2 respectively. It had a molecular formula of C_22_H_26_O_11_

The ligand CNP0166496 showed diverse interactions with multiple residues in the binding pocket of IDH1 and IDH2. It showed major H-bond interactions with Leu120, Lys126, Ile128 at a distance of 2.20 Å, 1.70 Å and 1.99 Å respectively, with IDH1 and Gln(B)316, Asp(B)312, Gln(A)316, Val(A)294 and Asp(A)312 with IDH2, at a distance of 1.85 Å, 2.10 Å, 2.04 Å, 1.95 Å and 1.99 Å respectively.

### 3.5. ADME/T Analysis

CNP0166496 shows significant pharmacokinetic and pharmacological potential as an excellent drug candidate. Its high melting and boiling points indicate stability, and its hydration free energy suggests good solubility. With pKa values ensuring stability across pH levels, a dipole moment reflecting moderate polarity, and a TPSA indicating strong biological interactions, it balances structural complexity and adaptability. The compound’s lipophilicity supports effective membrane permeability, and compliance with Lipinski’s Rule of 5 underscores its drug-likeness. Despite minor synthetic and reactivity alerts, it remains viable. Moderate water solubility and Caco-2 permeability are offset by reasonable human intestinal and oral absorption. As a P-glycoprotein substrate, it shows extensive tissue distribution and substantial plasma protein binding. Low BBB and CNS permeability reduce CNS side effects. Lacking major CYP enzyme interactions minimizes drug-drug interactions, and efficient clearance prevents accumulation. Its manageable safety profile includes predicted GHS class 5 toxicity, moderate AMES toxicity, no hERG inhibition, moderate oral rat toxicity, and no skin sensitization, DILI, carcinogenicity, or eye and respiratory toxicity. A low FDA recommended daily dose suggests high potency. Overall, the compound’s stability, favourable pharmacokinetics, and safety profile make it an excellent drug candidate.

### 3.6. Molecular Dynamics Simulation

#### 3.6.1. RMSD analysis

In Figure 5 (A), it is observed the backbone atoms of IDH proteins remained almost stable during the 100 ns simulation period for both the simulations. For complex 5(A), the backbone of the protein increased rapidly in the beginning till around 7-8ns with an RMSD of 6.5Å but later stabilized from till the end of the simulation indicating that the protein reaches a relatively stable conformation. This stability is crucial as it suggests that the protein maintains its structural integrity over the simulation period. The initial increase in RMSD is common as the system equilibrates, but the eventual stabilization indicates a good quality simulation. The ligand’s RMSD increases very gradually within the first 50 ns, reaching around 5Å, and fluctuates between 5-9 Å thereafter. This indicates that the ligand undergoes significant conformational changes or movements relative to the protein. The higher RMSD might suggest flexibility, which can be beneficial for the ligand to interact with different parts of the protein’s binding site.

**Figure 5.**
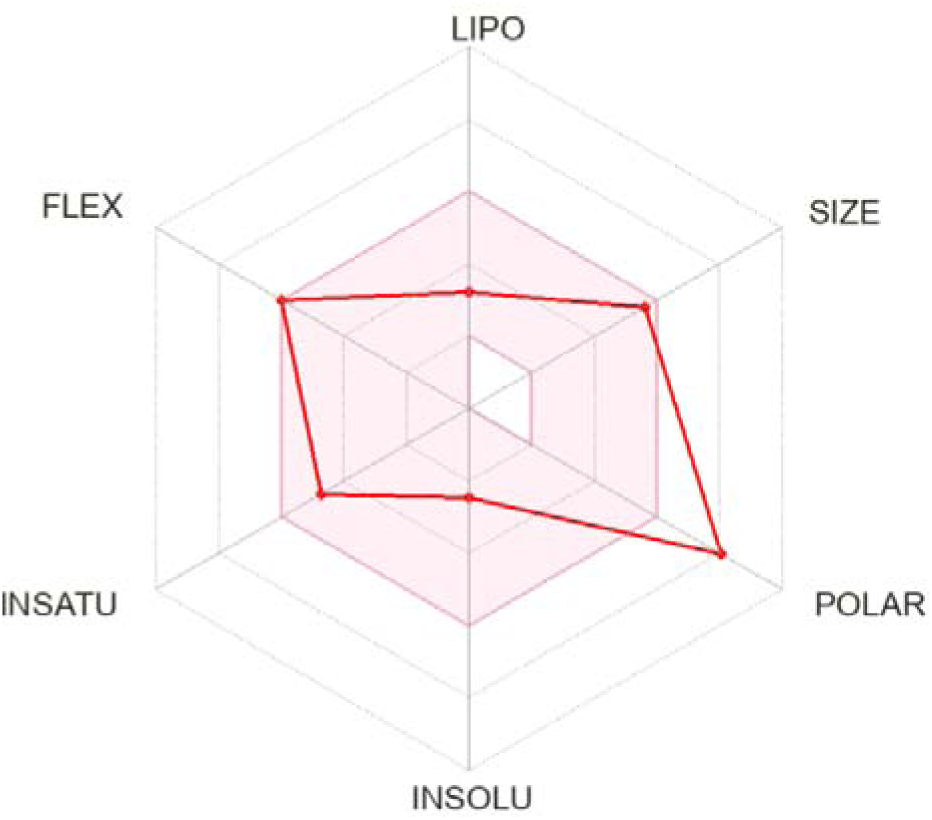
The bioavailability radar plot (obtained from SwissADME) depicting the excellent drug-likeliness of CNP0166496 (Ternstroside D). The reddish-brown area represents the optimal range for each property. The more detailed radar plot obtained from ADMETlab, are available in Figure S5

**Figure 6.**
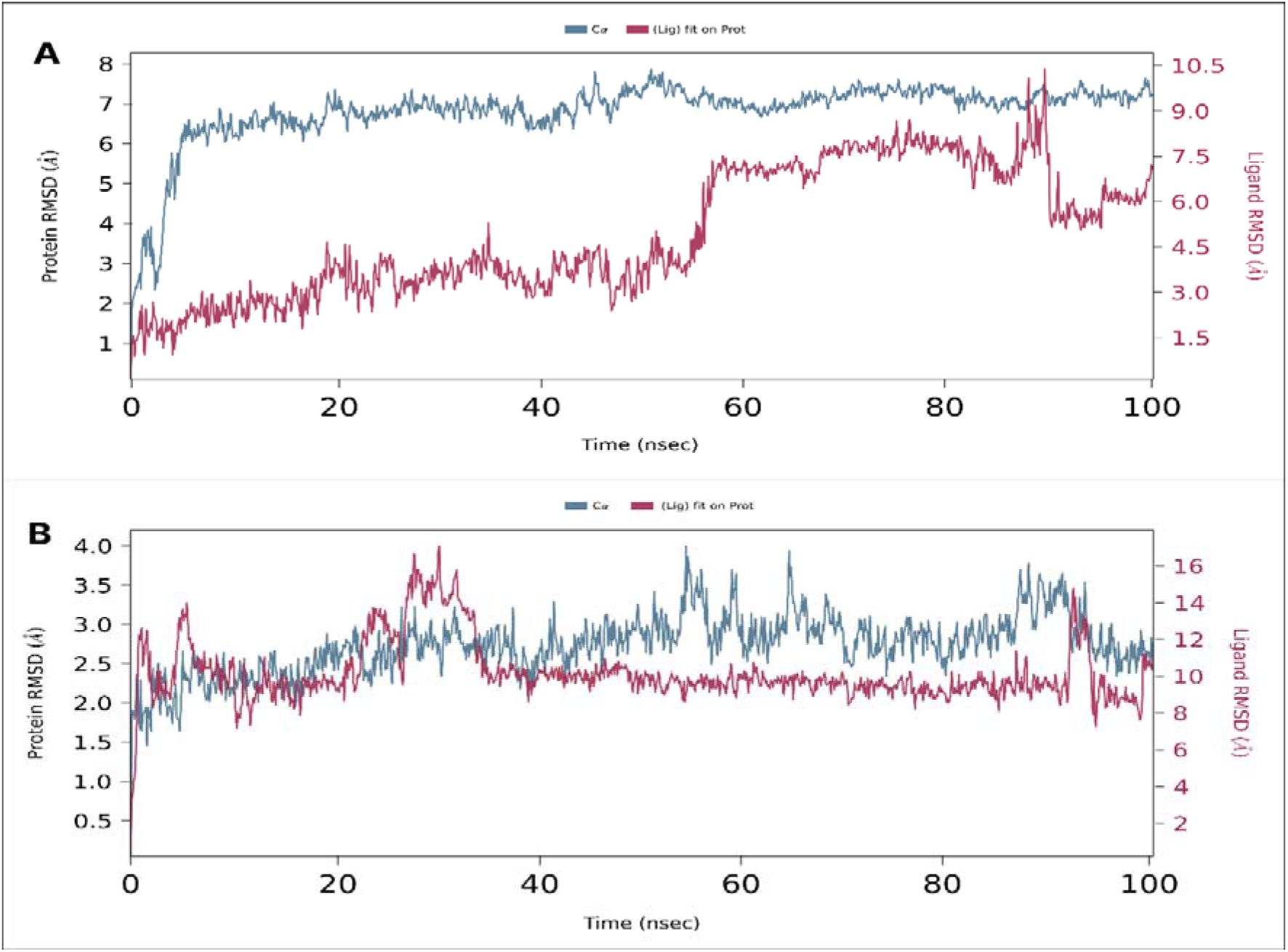
The line graph showing RMSD values of the complex structure extracted from ligand fit protein (ligand concerning protein) atoms. (A: IDH1 - CNP0166496 Complex; B: IDH2 - CNP0166496 Complex)

**Figure 7.**
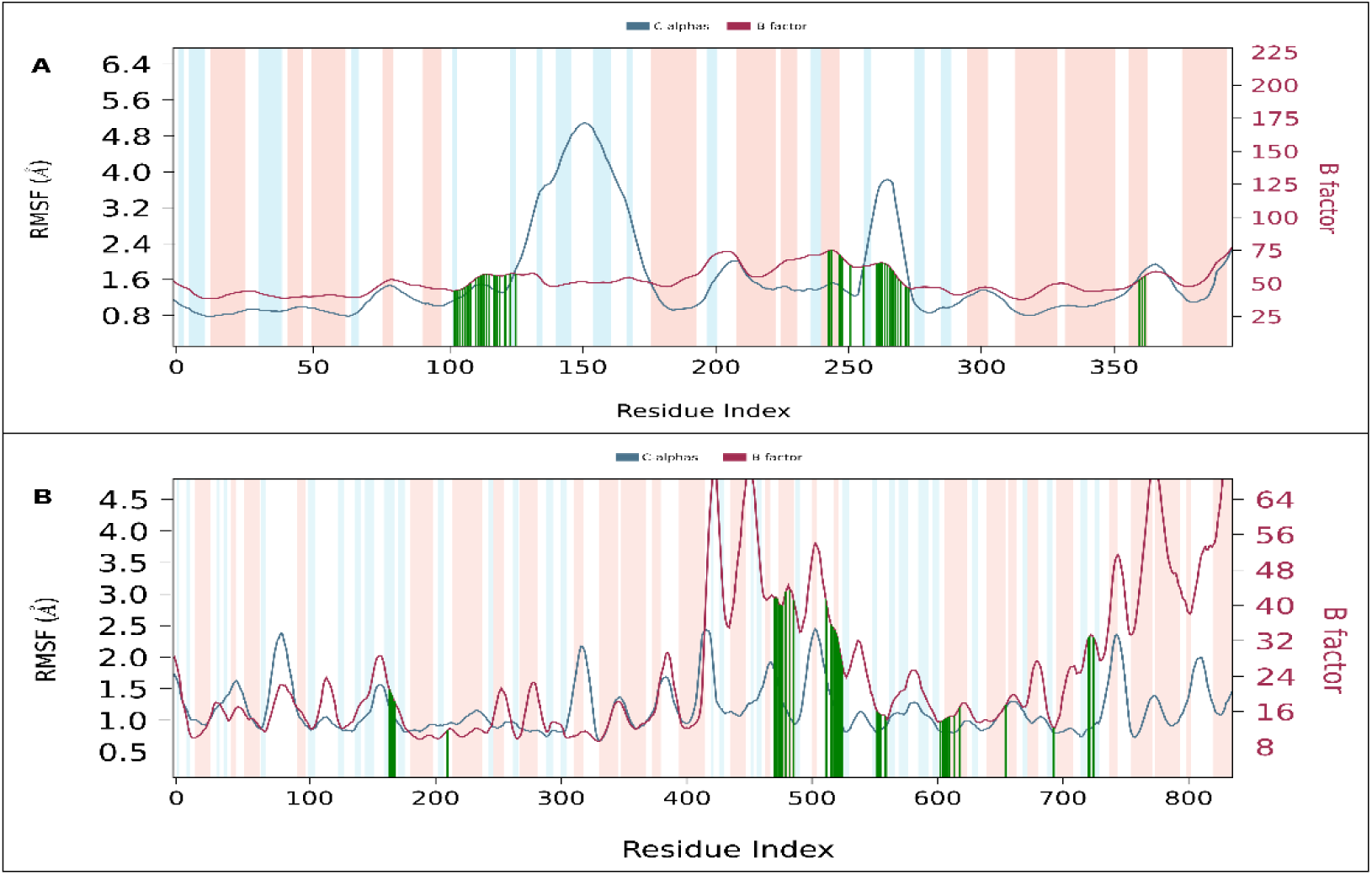
The line graph showing RMSF values of the complex structure extracted from protein residues backbone. The peaks of the blue line graph indicate the areas of the protein that fluctuate during the simulation. The maroon line graph indicates the B factor. The blue regions represent the alpha-helices and the orange region indicates the beta-helices. Protein residues that interact with the ligand are shown in green-coloured vertical lines. (A: IDH1 - CNP0166496 Complex; B: IDH2 - CNP0166496 Complex)

In Figure 5(B), The protein’s RMSD increases up to around 4Å at around 60ns. But overall, the protein remains quite stable from the beginning of the simulation. This consistent stability is a positive indicator of the reliability of the simulation and the potential for the ligand to interact effectively without destabilizing the protein. The ligand’s RMSD shows multiple fluctuations in the first 35-40 ns, but later it extremely stabilizes with a value of around 8.5Å. This relative stability suggests that the ligand maintains a consistent binding mode, which is crucial for effective drug design as it indicates strong and stable interactions with both the protein’s binding pocket.

#### 3.6.2. RMSF Analysis

In Graph A, the RMSF analysis of C-alpha residues reveals a dynamic protein structure with RMSF values ranging from 0.8 Å to 5 Å, indicating varying degrees of flexibility across different regions. The simulation captures significant peaks around residues 150 and 250. These peaks correlate well with B-factor fluctuations (25 to 225), affirming the simulation’s fidelity in capturing dynamic protein-ligand interactions.

Graph B displays a similarly robust RMSF profile, ranging from 0.5 Å to 2.5 Å, showcasing a well-defined protein-ligand interaction landscape. Notable peaks at residues 350, 500, 700, and 800 highlight distinct regions of flexibility, aligning closely with corresponding B-factor variations (8 to 64). This coherence between RMSF and B-factor data validates the simulation’s accuracy in depicting protein dynamics under ligand influence.

#### 3.6.3. Protein-ligand contact dynamics

In the present study, it is identified four types of interactions within MD simulation: hydrogen bonds, hydrophobic interactions, water bridges, and ionic bonds. The timeline interactions of the residues of the protein with the ligands are shown in Figure 9, which shows a stable interaction.

As in Figure 8A and 9A, the residues of IDH1(PDB: 5DE1) that commonly interacted with the co-crystalized inhibitor GSK321A and to CNP0166496 are Lys126, Tyr285, Trp124, Val281, and Gly284. On the other hand, in figure 8B and 9B, IDH2 also interacted in the same binding pocket in nearby residues to that of the co-crystallized inhibitor AG-221 (Enasidenib) like: B:Lys180, B:Gly145, A:Try210, B:His173, B:Ala174, B:His175, B:Lys133, B:Ser134, etc. indicating important interactions in the binding cavity

**Figure 8.**
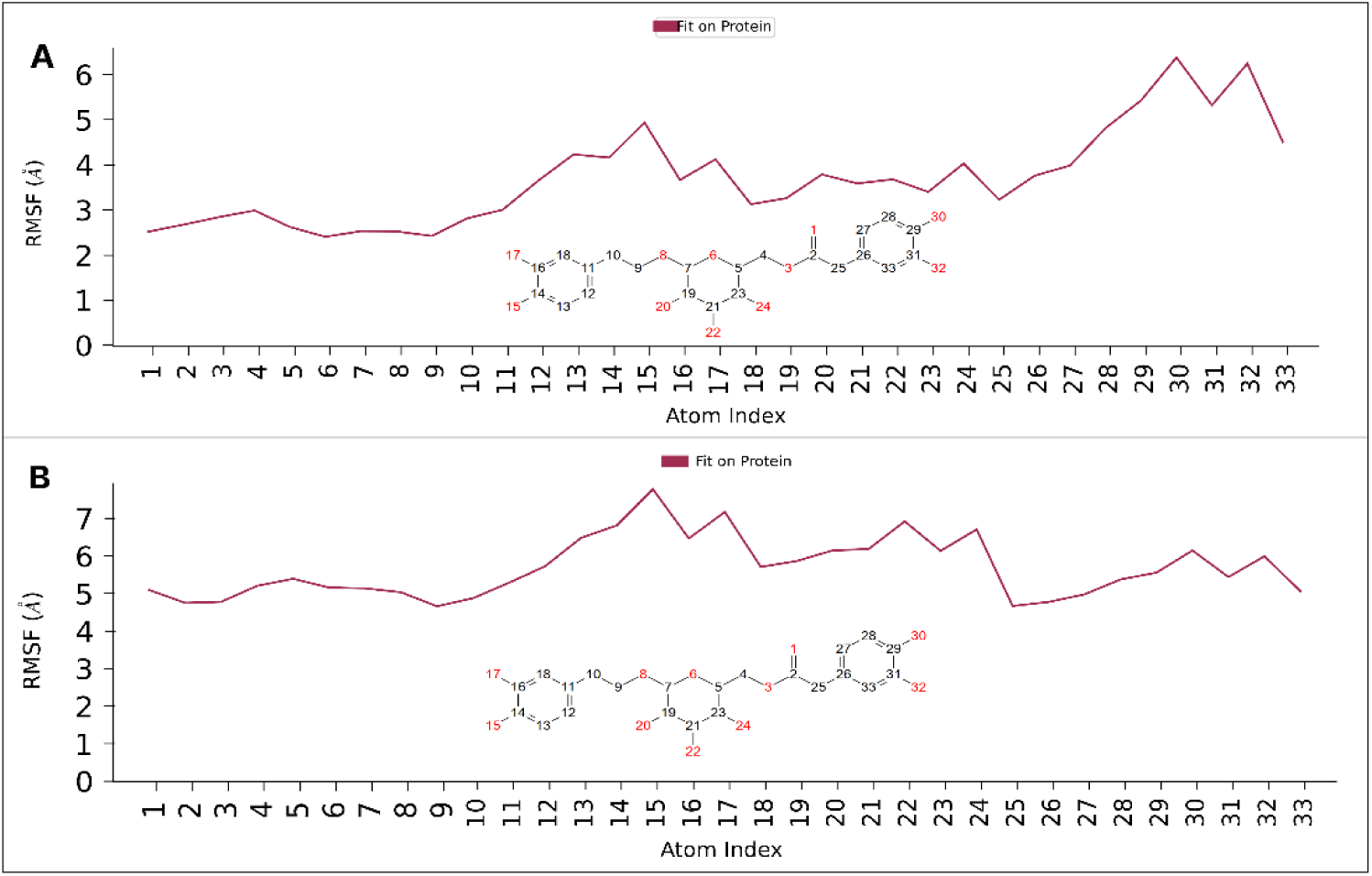
RMSF of Ligands’ atoms in the binding pocket of the IDH proteins (A: IDH1; B: IDH2)

**Figure 9.**
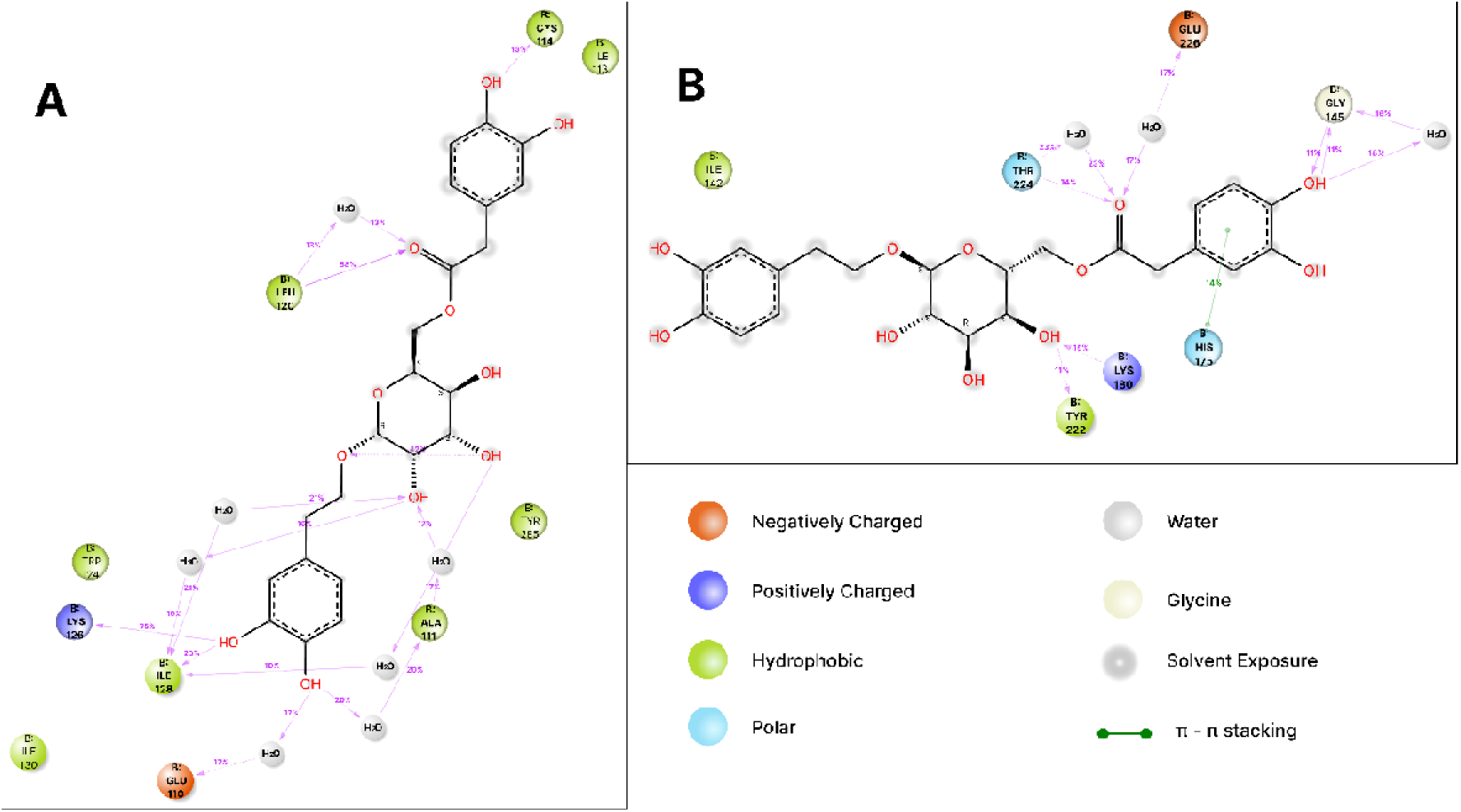
2D interaction diagram between protein-ligand complex (A: IDH1 - CNP0166496 Complex; B: IDH2 - CNP0166496 Complex)

#### 3.6.4. Properties analysis of ligand CNP0166496 (Ternstroside D

To validate the high stability of the examined complexes in an aqueous environment, we assessed the dynamic properties of ligand CNP0166496, which contribute to their stable interaction with the IDH1 and IDH2 proteins’ active sites. Figure S1(A, B) summarizes the six examined properties: Ligand RMSD, Radius of Gyration (rGyr), Molecular Surface Area (MolSA), Intramolecular Hydrogen Bonds (intraHB), Solvent Accessible Surface Area (SASA), and Polar Surface Area (PSA).

From Figure S1 (A, B), it is observed that the RMSD values of ligand CNP0166496 in complex with IDH1, remained generally stable with an average of 1.62 Å. The RMSD values ranged between 0.26Å at 10.6 ns and 3.22Å at 97.7ns. On the other hand, the RMSD values of the ligand with IDH2 showed an average of 2.34Å throughout the 100 ns simulation, varying between 1.32Å at 57.9 ns and 3.42Å at 94.1 ns. The slight variations in RMSD values for all the two ligands (<∼3Å) during the 100 ns MD simulation indicate the high stability of ligands CNP0375130 and CNP0256178 in the protein pocket. The average stability of rGyr values during the 100 ns MD simulation for CNP0166496 are 4.74Å and 4.90Å, respectively for IDH1 and IDH2. For the properties MolSA, SASA, and PSA of ligand, the values were observed in the range of (337-417Å and 335-426Å), (58-269Å and 191-536Å), and (289-305Å and 285-395Å), with averages of 389.36 Å, 140.99 Å, 340.70 Å and 400.77 Å, 290.47 Å, 368.90 Å respectively for complex with IDH1 and IDH2 respectively. These findings further support the stability of ligands CNP0166496 in the proteins’ environment.

### 3.7. Principal Component Analysis (PCA) and Dynamical Cross-Correlation Matrix (DCCM) Analysis

The standard PCA analysis (see Figure 10) of the ligand-bound IDH1 and IDH2 complexes offers insights into their conformational space and dynamic behaviour. Graph A (IDH1 bound to CNP0166496) shows data points clustered around the origin (PC1 ≈ 0, PC2 ≈ 2), indicating stable conformations and flexibility along PC1 (−2 to 12). This spread indicates that while the protein predominantly remains in a stable state, it explores various conformations, potentially enhancing its binding affinity and interaction with the ligand.

**Figure 10.**
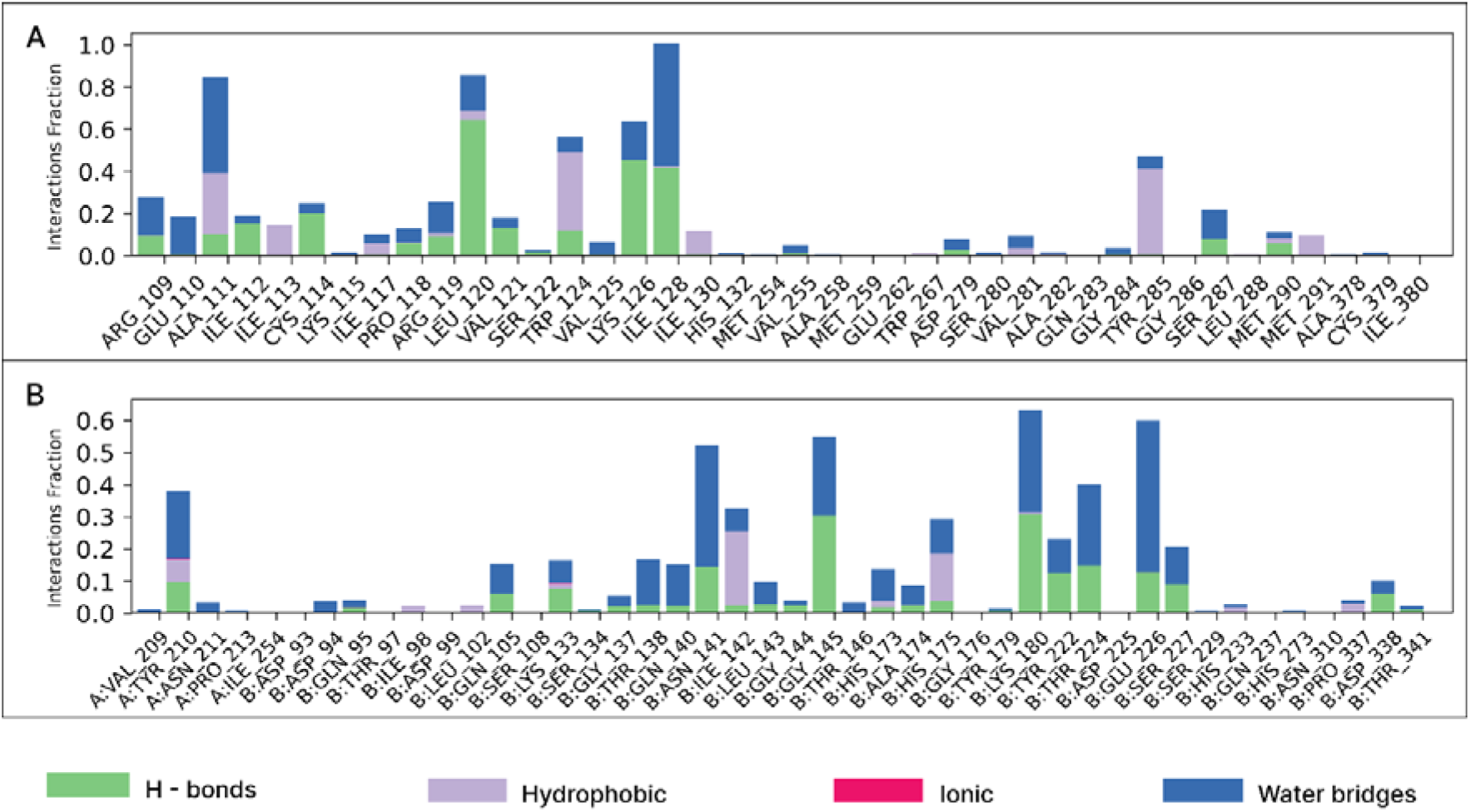
Interaction fraction histogram between protein-ligand complex (A: IDH1 - CNP0166496 Complex; B: IDH2 - CNP0166496 Complex)

Graph B (IDH2 bound to CNP0166496) exhibits dispersed data points across PCs, suggesting a broader conformational space and dynamic behaviour. IDH2 lacks a dense central cluster, indicating significant conformational changes upon ligand binding, which might be essential for its functional regulation. The Internal PCA, Multi-Dimensional Scaling (MDS), and t-distributed Stochastic Neighbour Embedding (t-SNE) graphs are shown in figure S5.

DCCM analysis (see Figure 11) of CNP0166496-bound IDH1 and IDH2 complexes reveals key insights into correlated motions induced by ligand binding. In IDH1 (Graph A), significant positive (red) and negative (blue) correlations are observed, particularly around residues 0-100, ∼140-170, and 300-400, indicating coordinated movements stabilizing the protein structure upon ligand binding. These motions likely play a critical role in the ligand’s inhibitory mechanism.

**Figure 11.**
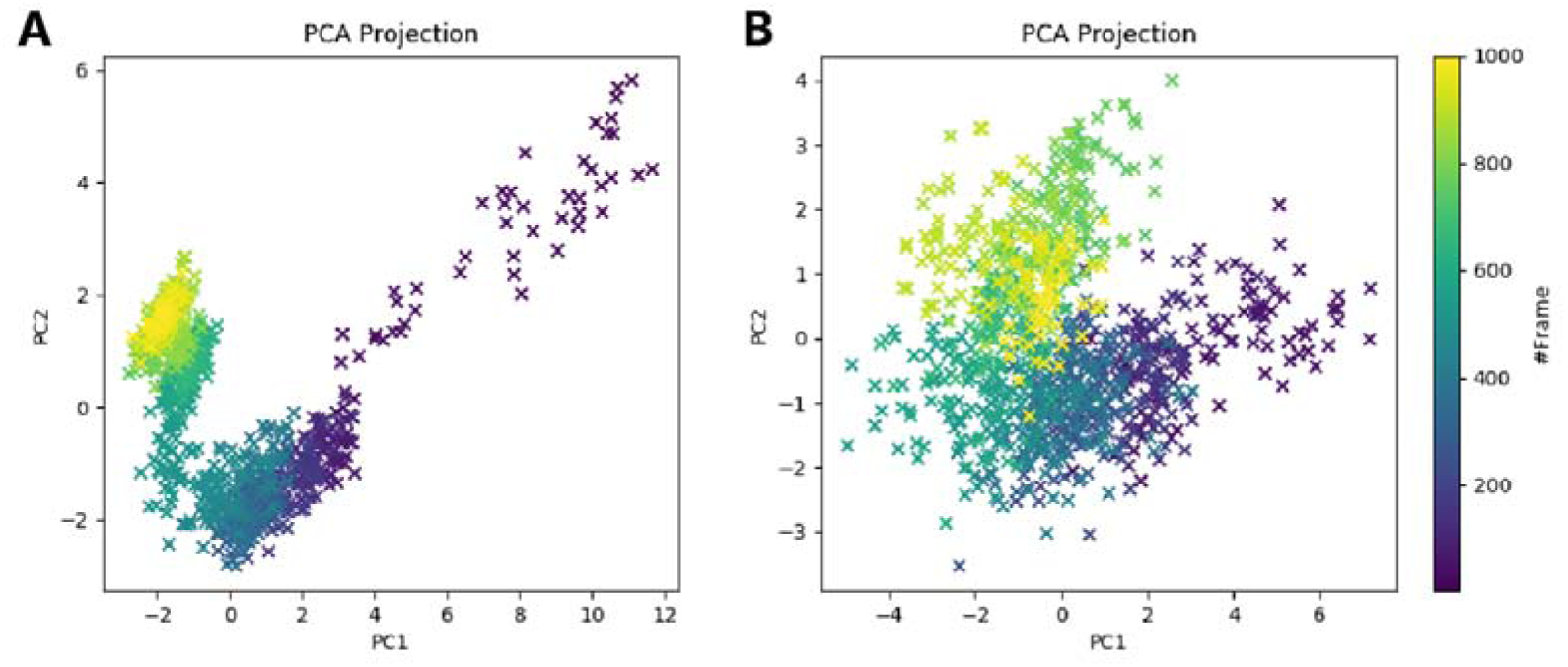
Standard Principal Component Analysis (PCA) graph of the trajectories of the two complexes (A: IDH1 - CNP0166496 Complex; B: IDH2 - CNP0166496 Complex)

**Figure 12.**
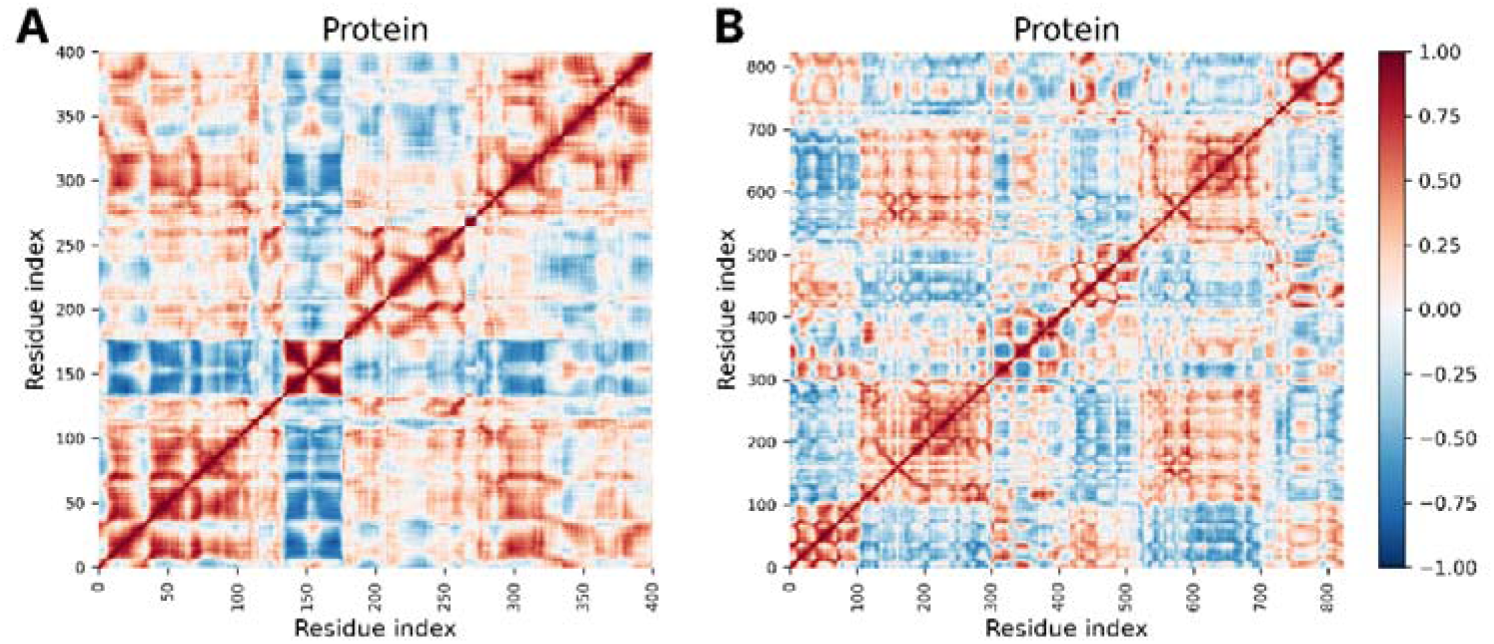
Dynamical Cross-Correlation Matrix (DCCM) Analysis graph of the trajectories of the two complexes (A: IDH1 - CNP0166496 Complex; B: IDH2 - CNP0166496 Complex). Red and blue colors indicate positive and negative correlations between residue pairs, respectively, while white represents minimal correlation. Strong diagonal dominance suggests local structural stability, with off-diagonal clusters indicating long-range interactions.

**Figure 13.**
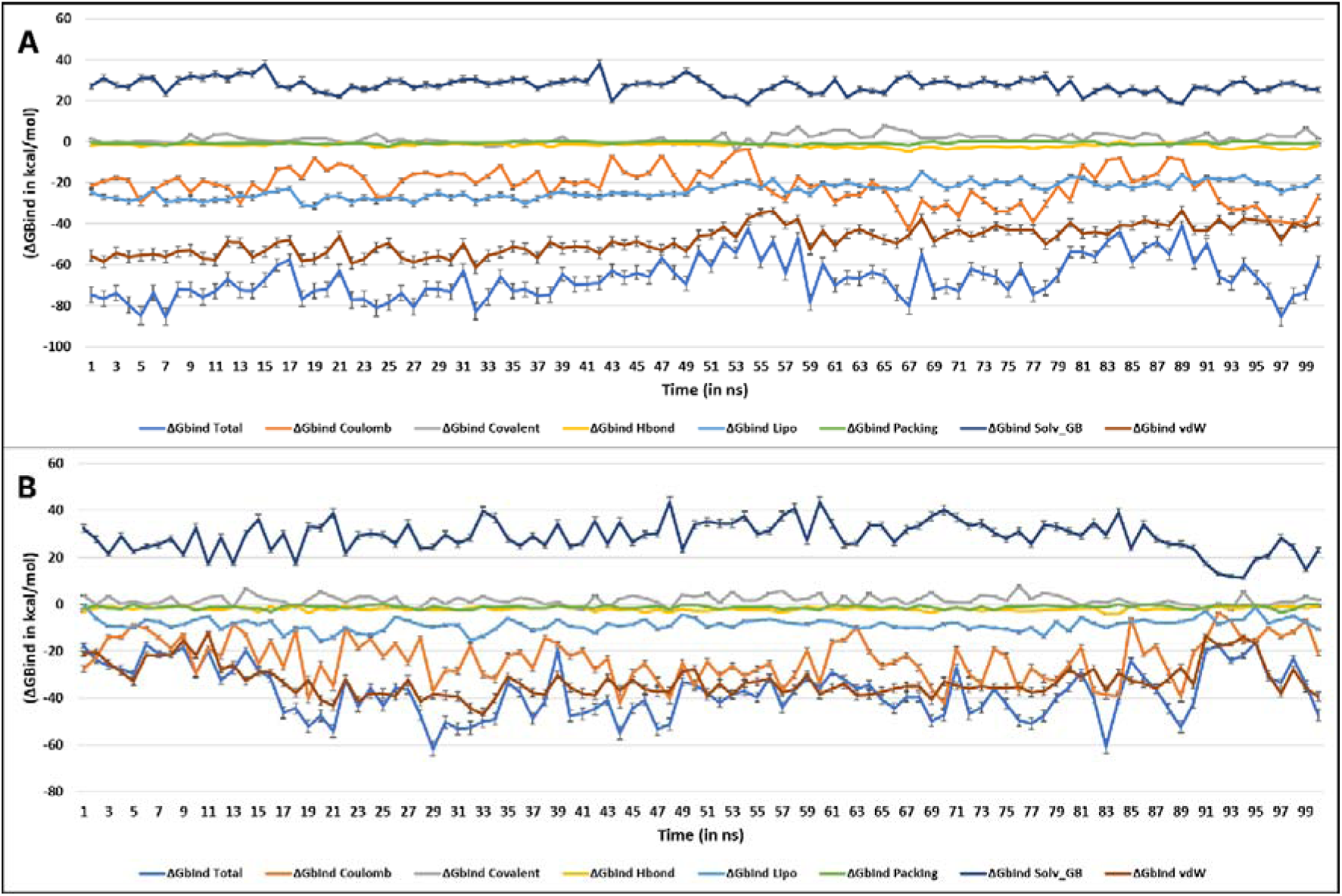
Post simulation MM-GBSA based Binding free energy (ΔG_Bind_ in kcal/mol) for (A) IDH1-CNP0166496 Complex; (B) IDH2-CNP0166496 Complex

IDH2 (Graph B) exhibits a more complex pattern with dispersed positive correlations and strong anti-correlations, particularly evident in the upper right quadrant. This suggests that CNP0166496 induces dynamic changes in IDH2, potentially leading to altered conformational states, which correlates with the PCA results. Correlated motions are notable around residues 100-300 and 500-700, indicating unique interactions and stabilization effects distinct from IDH1.

### 3.8. MM-GBSA Analysis

The parameters of free binding energies between the ligands were evaluated by the MM-GBSA simulations, which is the final step for checking the stability levels of the examined systems in the aqueous environment. The average values of calculated MM-GBSA energy parameters are presented in Table S4, containing binding free energies, Coulomb energy (Coulomb), Covalent bonding (Covalent), Hydrogen bonding (H-bond), lipophilic bonding (Lipo), π-π stacking interaction (Packing), solvent generalized bonding (SolvGB), and Van der Waals bonding energy (VDW) over the 100ns time frame.

From Table S5 and S6, the ΔG_bind_ Total of IDH1-CNP0166496 Complex had an average value of −66.66 kcal/mol, maximum value of −104.54 kcal/mol, minimum of −40.85 kcal/mol, and with a standard deviation (SD) of 9.96. The majority of it was contributed by ΔG_bind_ Coulomb (−21.49 kcal/mol), ΔG_bind_ Lipo (−23.78 kcal/mol) and, ΔG_bind_ vdW (−48.05 kcal/mol). The average ΔG_bind_ Total shown by the IDH2-CNP0166496 complex was −37.46 kcal/mol having a maximum value of −104.54 kcal/mol, minimum of −40.85 kcal/mol, and with a standard deviation (SD) of 10.82. The majority of it was contributed by ΔG_bind_ Coulomb (−29.23 kcal/mol), ΔG_bind_ Lipo (−8.75 kcal/mol) and, ΔG_bind_ vdW (−32.61 kcal/mol).

## 4. DISCUSSION

This *in silico* study aimed to identify a dual inhibitor targeting IDH1 and IDH2 mutations implicated in AML. Virtual screening identified a lead compound, CNP0166496 (Ternstroside D), exhibiting high docking scores and binding affinities with both IDH1 (−14.2 kcal/mol) and IDH2 (−16.822 kcal/mol) as predicted by MM-GBSA calculations, which are significantly higher than those of the native inhibitors GSK321A for IDH1 (−9.6 kcal/mol) and Enasidenib (AG-221) for IDH2 (−8.9 kcal/mol) (Bello *et al*., 2023). According to the COCONUT database, Ternstroside D was retrieved from various database collections, including NP-MRD, ChEBI, SuperNatural2, UNPD, and NPASS, highlighting its well-documented presence in natural product databases. Ternstroside D is a phytocompound classified as a triterpenoid saponin, characterized by a steroid-like core structure with glycosidic moieties attached, and is isolated from plants in the genus *Ternstroemia* within the Theaceae family. Notably, *Ternstroemia gymnanthera* and *Ternstroemia japonica*, native to Asia and East Asia respectively, are known sources of this compound. Ternstroside D is a β-D-glucoside featuring a 2-(3,4-dihydroxyphenyl)ethoxy residue at the anomeric position and a [(3,4-dihydroxyphenyl)acetyl]oxy residue at position 6. Notably, its antioxidant activity has been validated *in vitro* (Jo *et al*., 2006). The compound exhibited favourable pharmacokinetic and pharmacological properties, suggesting good drug-likeness according to Lipinski’s Ro5. It demonstrated acceptable water solubility, Caco-2 permeability, and limited blood-brain barrier penetration, indicating potential oral bioavailability and reduced central nervous system side effects. Minor synthetic challenges and potential reactivity issues could be addressed through medicinal chemistry optimization (Brown & Boström, 2016). RMSD analysis indicated stable protein-ligand complex formation throughout the 100 ns simulations for both IDH1 and IDH2. While the ligand showed some conformational flexibility, it maintained interactions with the binding pocket. The study identified interactions with residues like Lys126, Tyr285, Trp124, Val281, and Gly284 in IDH1, which are crucial for binding of established IDH1 inhibitors like GSK321A (Liu *et al*., 2023). This overlap suggests that Ternstroside D might utilize a similar binding mode, potentially mimicking the inhibitory mechanism of existing drugs. Similarly, interactions with residues like Lys180, Gly145, Try210, His173, and Ala174 in IDH2 were observed, resembling the binding mode of inhibitors like Enasidenib (AG-221) (Morell *et al*., 2022), suggesting potential competitive binding with the established inhibitor for the same active site. The presence of hydrogen bonds between Ternstroside D and both IDH1 and IDH2 suggests a crucial role in stabilizing the protein-ligand complex, which could be further analyzed to identify specific functional groups on the ligand contributing to binding affinity. Hydrophobic interactions likely play a role in anchoring the ligand within the active site, with understanding these interactions valuable for future structure-activity relationship (SAR) studies aimed at optimizing potency. The identified interactions with residues such as Lys126, Tyr285, Trp124, Val281, and Gly284 in IDH1 suggest a mechanism similar to established IDH1 inhibitors like GSK321A, while interactions with Lys180, Gly145, Try210, His173, and Ala174 in IDH2 resemble the binding mode of inhibitors like Enasidenib (AG-221), targeting the substrate binding pocket and likely competitively inhibiting the mutant enzyme’s ability to convert isocitrate to 2-HG. Future directions might involve evaluating the inhibitory activity of Ternstroside D against cell lines harbouring IDH1 and IDH2 mutations to validate the in silico predictions, investigating the mechanism by which Ternstroside D inhibits IDH enzyme activity, assessing the selectivity of Ternstroside D for mutant IDH proteins over other related proteins, and conducting structure-activity relationship (SAR) studies to optimize the structure of Ternstroside D to improve binding affinity and potency if possible.

## 5. CONCLUSION

Ternstroside D (CNP0166496), a natural product identified through a comprehensive computational pipeline, emerged as a promising dual inhibitor for both IDH1 and IDH2 mutations. The study’s robust methodology, encompassing pharmacophore modelling and screening, docking, and MD simulations, provides compelling evidence for Ternstroside D’s favourable binding affinity and stable interactions with the target proteins. Furthermore, *in silico* ADME/T analysis suggests its potential for successful drug development.

These findings pave the way for further exploration of Ternstroside D’s therapeutic potential. *In vitro* and *in vivo* studies are crucial next steps to validate the in-silico data and translate this discovery into a tangible clinical benefit for AML patients harbouring IDH mutations. Moreover, this study underscores the power of *in silico* methodologies as valuable tools for accelerating drug discovery, particularly in the realm of natural products with immense therapeutic potential.

**Table 1.** Top interacting residues of CNP0166496 (Ternstroside D) with the IDH proteins, XP docking score (kcal/mol) and MM-GBSA ΔG_bind_ energy (kcal/mol)

| Protein | H-bond Interacting Residues | Docking Score | MM-GBSA ($\Delta G_{bind}$) |
| --- | --- | --- | --- |
| IDH1 | L120, K126, I128 | -14.200 kcal/mol | -84.45 kcal/mol |
| IDH2 | Q(B)316, D(B)312, Q(A)316, V(A)294, D(A)312 | -16.822 kcal/mol | -60.73 kcal/mol |

